# CA1 and CA3 pyramidal cell morphologies under Alzheimer’s disease Amyloid beta interaction

**DOI:** 10.1101/187476

**Authors:** Yuri Elias Rodrigues, Josiane da Silva Freitas

**Affiliations:** Federal University of Rio Grande do Sul, Porto Alegre, Brazil; Lund University, Lund, Sweden

**Keywords:** Alzheimer’s disease, realistic morphology, glutamatergic synapses

## Abstract

Alzheimer’s disease (AD) is a fatal neurodegenerative disorder, which imposes a growing burden on society and health systems worldwide. Remarkable, computational models for AD are rare in comparison to epilepsy and other neuropathologies. Also, neuromorphological variability in the available models is usually neglected. Here, we evaluated geometrically detailed CA1 and CA3 pyramidal neurons from rat’s hippocampus and its firing probability face to Amyloid-β (Aβ), a pre-plaque soluble confirmation. Such comparison is invaluable since the hippocampus acts as a structural predictor of AD progression. A stimulation protocol designed to elicit a single spike is used to access neuronal vulnerability against the Aβ oligomers. Our experiment shows that under the same conditions there is a firing facilitation for CA3 neurons in comparison to CA1.

## 1 Introduction

Alzheimer’s disease (AD) is a neurodegenerative disorder, currently irreversible, characterized by memory impairment and affecting mainly elderly people [1]. Despite that, there is a broad range of therapeutics strategies to retard its progression, from cognitive enhancement training [2] to the cholinergic modulation [3]. Unfortunately, such approaches act only as palliative care for mild and advanced late onset AD [4]. For this reason, early-stage identification and computational modeling of AD is useful for disease-altering status and drug discovery research [5]. A key region for understanding AD progression is the hippocampus which its shrinking usually leads to memory and spatial coordination loss [5]. Computational models for AD objectively measuring hippocampal alterations are mainly focused on synaptic dysfunction [6] and proteins aggregation dynamics [7]. Yet, they cover only partially the AD neuropathology aspects when proposing drug targets [7]. For instance, in [6] it is proposed an ion-channel conductance modification to reduce AD effects mediated by Amyloid-β (Aβ) protein based on a single CA1 neuron. Here, it is considered a set of reconstructions of CA1 and CA3 pyramidal neurons (glutamatergic) and its spiking probability face to an Aβ implementation [6]. The Aβ cascade is one of many hypothesis of AD progression and supports the amyloid Aβ oligomeric build-up as the main driver of neurodegeneration [8]. Our results allows one to evaluate region-based vulnerability and how different neuron geometry resists to Aβ exposure.
The hippocampus has been received attention due to its role in navigation, neurodegeneration and memory processes [9]. Thus, neuropathology simulation in its regions, e.g. CA1 and CA3, is of great interest. The CA1 region is credited to participate in new memory formation and memory strengthens [10]. Whereas CA3 represents the major component of CA1 activity through Schaffer collaterals pathway [3]. Individual regions contributions for different hippocampal functions has been explored by computational studies which are publicly available at ModelDB repository. However, region differential dysfunction lacks to be researched for AD since its mechanisms are not completely known [1]. Advancement in AD neuropathology modeling remains at least limited by how well documented the damaged components. For instance, the CA1 region which shows to be highly affected by tau protein toxic build-up in early stages of AD has scarce tau’s electrophysiology measurements suitable for virtualization [11] [4].

Computational neuroscience allows simulates brain dynamics in several scales opening new possibilities of experiments. There are multiple directions in which computational models would propose a way to change the AD disease status at least in cellular and molecular level. The number of assumptions [7] required to cover AD experimental findings [8] is a limitation for simulation neuroscience since there is no consensus regarding its initial causes [12]. This scenario is worsened by the lack of connection between distinct methods with distinct temporal and spatial resolution to observe neuropathologies [13], e.g. neuroimaging and electrophysiology. A neurocentric strategy for therapeutics discovery is to adjust the firing properties of AD-affected neurons to behave as healthy ones. Experimentally, a promising research by [14] shows AD neuronal oscillations induced by optogenetics enables stimulates a healthy spiking patterns, increasing Aβ endocytosis. Computationally, focusing on the single neuron scale, a drug target is identified by ion-channel modifications in a detailed CA1’s pyramidal cell model in order to mitigate Aβ effects in the spiking probability [6]. However, in such approach, there are no considerations regarding how morphological variations from the same and other hippocampal regions would affect the spike probability. Here let’s observe the reaction of CA1 and CA3 neuromorphologic strains when the Aβ is included through conductance reduction in the synaptic transmission for a compartment neuron model.

## 2 Data and Methods

The simulations were carried out in Python (v 2.7) and NEURON (v 7.2) [15] and the neuromorphologic reconstruction were obtained online from Neuromorpho.org repository. Rat’s hippocampus CA1 (n=41) and CA3 (n=43) neurons were filtered by complete basal and apical dendritic tree, incomplete axon and with soma. In order to evaluate the spike probability, which is defined as the chance of a given synaptic input to provoke a spike, a stimulation protocol is adapted from [6].

Here, it is used two sets of 25 AMPA-induced synapses distributed between the proximal and distal regions in apical dendrites. Apical tree receives synapses with 10 ms of time decay whereas proximal receives 5ms. We implemented the synaptic distribution rule by depending on the neuron size. The proximal apical region is considered until 3/4 of distance from soma to the neuron higher segment, whereas distal region is set after 3/4 [9]. Peak synaptic conductances are set to 0.87 nS for the proximal, whereas distal synapses are three times weaker. AD effects in neurons are stimulated through membrane conductances reductions randomly imputed for a given percentage of neuron surface area [6]. When a synapse is set in a segment affected by AD its peak synaptic conductance is halved [16]. The start time activation for synaptic stimulation is given by a Gaussian distribution (μ=50 ms, σ^2^=5 ms) to account for γ-cycle positive sweep [6]. Moreover, distal synapses are activated with a time delay of 5ms to simulate synaptic arriving delay between Perforant Path and Schaeffer Collaterals [17]. Despite this delay being biologically plausible for CA1, we keep the same stimulation protocol to CA3 in order to compare synaptic summation among the different regions. Using L-measures software [18] is possible to evaluate topological and compartment level measures for CA1 and CA3 morphologies as depicted in the Figures 1 and 2. The complete dataset of neuron reconstructions and its extracted features are available at https://github.com/yurier/CA1_CA3_morpho/ repository for further studies in pattern recognition.

**Fig. 1.**
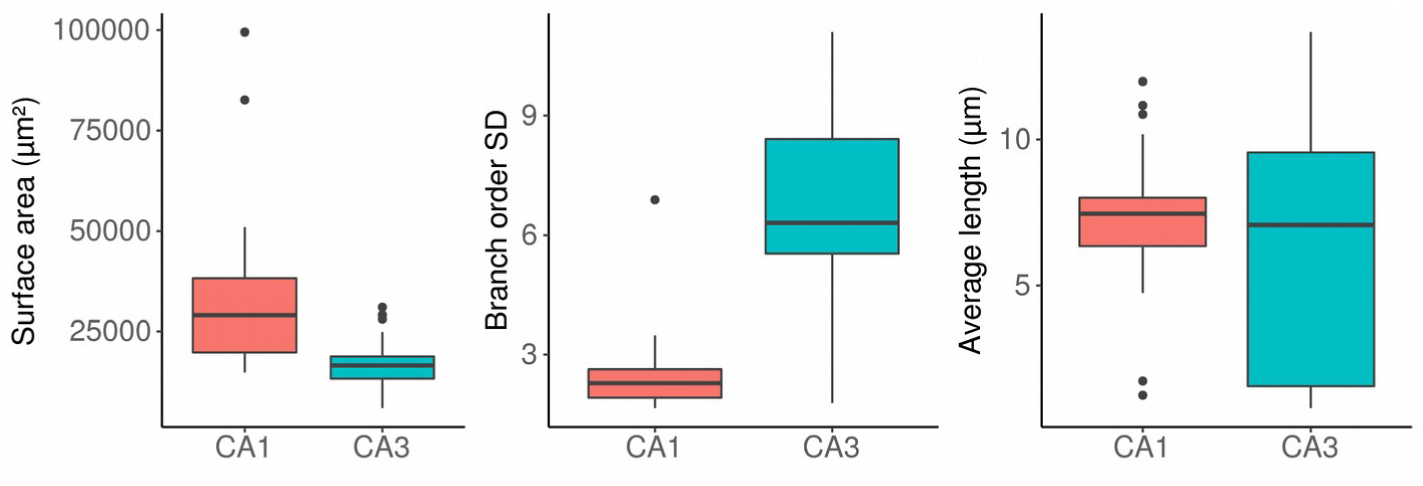
From left to right boxplots, respectively, the neurons surface area, the variability of branch order, and the average compartmental length.

**Fig. 2.**
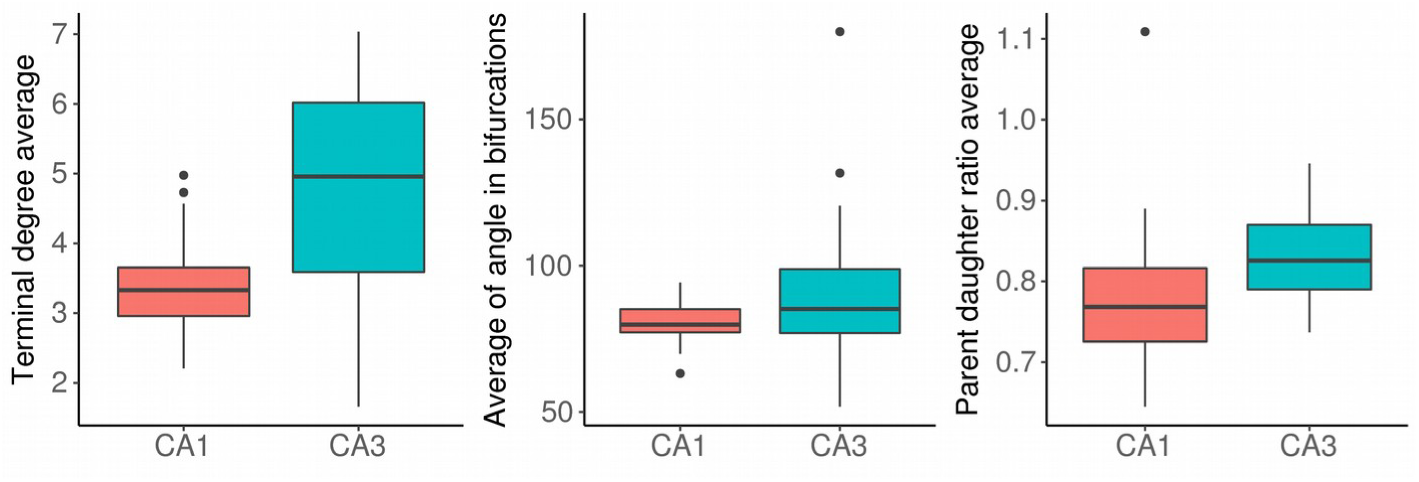
From left to right boxplots, respectively, the terminal degree average, the average angle in bifurcations, and the parent daughter ratio average.

Membrane resistance properties for batch simulation were fixed differently for CA1 and CA3 based on the description given [17] and [19], respectively. Each percentage of the affected membrane is repeated 10 times and averaged to measure the spike probability. The ion-channels density vary linearly with the distance from the soma as described in [17]. The CA1 implementation is equivalent for CA1 and CA3. The ion-channels used are Ih, KA, KDR and Na, the conductances reductions to resemble Aβ effect are respectively, KA (-60%), KDR (-40%) and Na (-50%) as done in [6].

## 3 Results and Discussion

Computational neuroscience applied to neuropathologies provides drug-targets to be tested in order to simulate healthy neuron behavior [7]. However, AD computational models have been quite a few in comparison to other neuropathologies [7]. Approaches to AD by means of computational modeling explores different levels from molecular to microcircuitry looking forward to possible interventions to reduce AD effects [7]. An AD model previously proposed by [6] suggests a drug-target based on a single pyramidal CA1 neuron morphology, however disregarding between-region variability in favor to a detailed ion-channel investigation. The fine-tuning for drugs obtained in silico environment should account for as much as possible target variability in order to avoid non-desired interactions. Here, an AD computational model of Aβ effect is simulated using multiple neuromorphologies from hippocampus CA1 and CA3 offering a background for future intervention studies [20]. Also, it is shown how realistic morphologies are impaired differently through ion-channels and synaptic transmission defects.

Single neuron experiments considering neurons as isolated units are useful to understand its computational roles in an environment without untraceable interactions as complex networks. Such paradigm allows evaluating how different neuron shapes contribute to the firing probability [21]. Morphology is one of the many determinants of spiking profile. Other features such as characteristic plasticity and wiring connectivity patterns would give rise to different firing patterns. An example of that is given by [21] which show the dominance of spike probability between CA1 and CA3 depends on the brain state activity (e.g. theta oscillations periods and rapid eyes movement) and the firing pattern (e.g. burst firing and single spike). They show that CA3 pyramidal cells have a higher probability of long bursts. Conversely, for the states studied by [21], CA1 presents an overall higher firing rate. Here, using the defined synaptic stimulation protocol, the morphology-based comparison suggests that CA3 neurons have a higher average probability in single spike firing in comparison to CA1. The same dominance is found given when using the implementation of membrane proportion affected by the Aβ [6].

It is noticeable by the Figure 3, despite CA1 has a basal firing probability lower than CA3 under same inputs, the CA3 has faster decreasing when Aβ membrane affected grows. Currently, there are no studies defining the minimum firing activity in terms of population spike probability to sustain an expected behavior for a given task in CA1 and CA3. That is, the maximum damage a region can tolerate prior to its multiple related functions became impaired is unknown. Another difficulty is how to measure this damage. Supporting evidence suggest that CA1 has greater loss of neuronal density in comparison to CA3 than any other hippocampal area in humans [22]. However, research is required in order to define the most vulnerable function and its rate of impairment in a scale that allows the computational modeling, e.g. neuromorphologic statistics, wiring connectivity patterns, ion-channel density, glial depletion, excitatory-inhibitory imbalance.

**Fig. 3.**
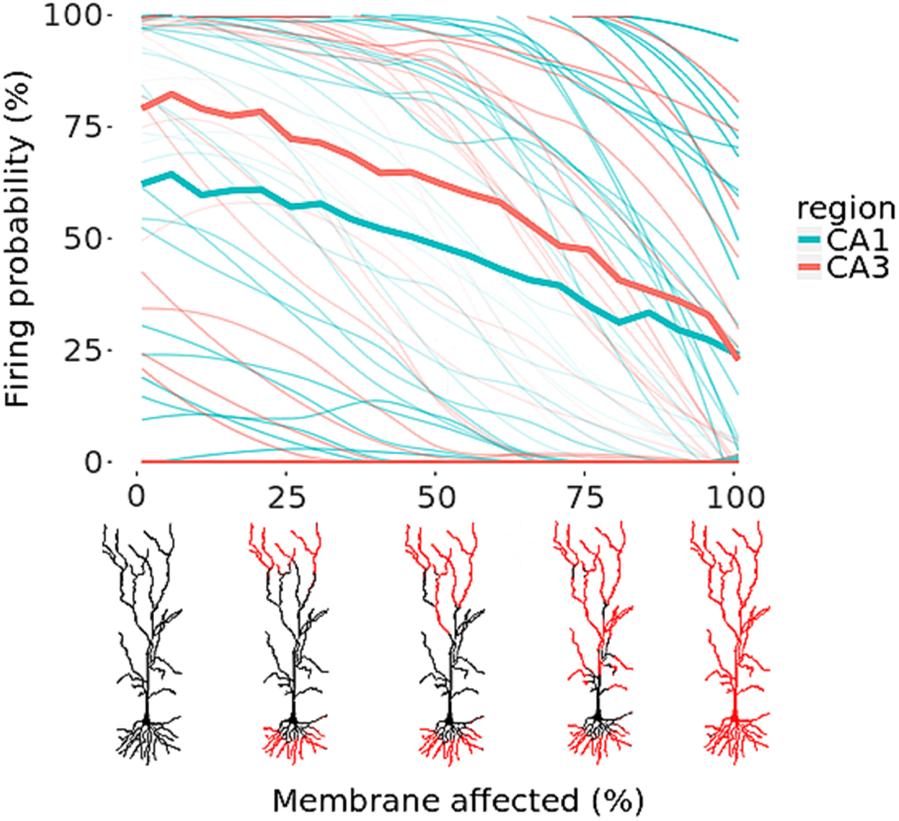
Firing probability average for CA1 and CA3 regions given the Aβ in the membrane. The smoothed thinner lines are the individual firing probabilities.

Regarding Figure 3 results, one would observe that under the fixed stimulation protocol there are similarities in firing probability between CA1 and CA3. For instance, some neurons in both regions despite having all compartments affected by Aβ they are able yet to elicit a spike. Such feature is more present in CA1 than CA3, respectively 7 and 2 neurons present this behavior. Conversely, the stimulation protocol was too weak for a group of 11 CA1 neurons, whereas all CA3 neurons have their average probability higher than zero.

Detailed morphology is crucial here since they incorporate dendritic computation and synaptic integration profiles which are unique neuron properties. This is relevant to extend this implementation to age-related dendritic defects which play an important role in AD pathology. The bottleneck of detailed geometries is the longer processing time. Alternatively, the ball-and-stick neuron model would present a higher performance gain for network simulation. However, in order to implement for networks, the Aβ mechanism presented here the membrane mechanisms which control affected area should be parametrized. There are efforts in modeling networks using a wide range of AD features [7]. For instance, in [23] it is implemented a network model by fitting AD synaptic loss parameters obtained from neuronal culture exposed to Aβ. Another network example is given by [24] in which the synaptic loss is simulated conceptually using Venn’s networks. Networks approach focusing on self-sustained activity would be the suitable computational paradigm to understand the memory dysfunction in AD and its dynamics in a future.

## 4 Conclusion

Here, one of the main hypothesis of AD, the Aβ protein accumulation, is simulated in order to demonstrate the role of morphologic features in synaptic integration. There are differences in the way which different cell types respond to Amyloid-β (Aβ) suggesting that morphologic variability must be accounted for drug-targeting discovery. To improve biological plausibility more AD features should be revisited. For instance, the substantial loss of dendritic tree complexity, the dynamic neuron branching, and the neurofibrilary tangles exposition. Such advancements are constrained by data resources which are harder to sample than neuron morphology reconstructions. AD and computation neuroscience would benefit by how neurological disorders are modeled in order to understand the disease or even to find a way to give AD patients a way to live better. Maybe a future in which we cure AD is bounded to the way in which data is produced and shared. Due to initiatives like KnowledgeSpace by Human Brain Project and the NeuroCurator great tools are available to help neuroscientists in their goals. Is important to strengthen that even with data sharing being a consolidated policy there is a significant appeal from advancement in techniques and theory.

